# Millennium-old Pathogenic Mendelian Mutation Discovery for Multiple Osteochondromas from a Gaelic Medieval Graveyard

**DOI:** 10.1101/2022.02.02.478802

**Authors:** Iseult Jackson, Valeria Mattiangeli, Lara M Cassidy, Eileen Murphy, Daniel G Bradley

## Abstract

Only a limited number of genetic diseases are diagnosable in archaeological individuals and none have had causal mutations identified in genome-wide screens. Two individuals from the Gaelic Irish Medieval burial ground of Ballyhanna, Co. Donegal, showed evidence of bone tumors consistent with the autosomal dominant condition multiple osteochondromas. Genome sequencing of the earlier individual uncovered a missense mutation in the second exon of *EXT1*, a specific lesion that has been identified in several modern patients. The later individual lacked this but displayed a novel frameshift mutation leading to a premature stop codon and loss of function in the same gene. These molecular confirmations of a paleopathological diagnosis within a single rural ancient context are surprisingly disjunct, given the observation of clusters of this disease in modern isolated populations and a *de novo* mutation rate of only 10%.

## Main Text

Cases of Mendelian genetic disease have been uncovered in the archaeological record, but diagnosis is restricted to osteologically visible conditions, with more commonly encountered conditions including achondroplasia, multiple epiphyseal dysplasia and Léri-Weill dyschondrosteosis ^1^.

Our study focuses on two adult male individuals (Sk197 and Sk331) buried in a Medieval graveyard at Ballyhanna, near the town of Ballyshannon in Co. Donegal, Ireland. Each displayed multiple bony tumors suggestive of multiple osteochondromas (MO), a rare, autosomal dominant bone condition^2^. Although these tumors are normally benign, the condition can result in limb deformity, reduced stature, compression of nerves, and, more rarely, malignancy^3^.

The rural burial ground at Ballyhanna was associated with a small Medieval church, constructed after the middle of the 13th century AD, although it is possible that an earlier wooden church was present on the site. The land at that time would have been owned by a bishop and the estate lands would have been managed by an *erenagh* (estate manager). Radiocarbon dating showed that the earliest burials dated to the late 7th to early 8th century, but the vast majority of individuals were interred between AD 1200 and 1650, when the area around Ballyshannon was under the autonomous control of the Ó Domnaill Gaelic lords. As such, the graveyard at Ballyhanna can be considered to have essentially contained the remains of a Gaelic Medieval population. Those buried at Ballyhanna would have comprised the lower classes and included tenant farmers, labourers, merchants, artisans, clergy and the very poor^4^.

Radiocarbon dating revealed that the two individuals were definitely not contemporaneous and were potentially separated by several hundred years. Of the two, Sk331 lived more recently (dated AD 1031-1260; UBA-11442) (**Figure S1; Table S1**) and was the more severe case. He displayed extensive bilateral osteochondromas, both sessile and pedunculated in form, on most bones throughout his skeleton (**Figure 1b.**). He also had a short stature compared to other adult males at Ballyhanna (158.3 cm), displayed a major deformity of his left forearm due to shortening of the ulna (Type 1^5^), had unequal bone lengths due to the lesions, as well as a range of orthopaedic deformities that affected his hips, knees and left ankle; all of which are consistent with this condition^6^. He died as a young adult (18-25 years). Sk197 was dated to AD 689-975 (UBA-11443) and was slightly older (30-40 years) when he died. While multiple osteochondromas were evident throughout his skeleton, they were generally less pronounced than those evident in Sk331. Limb length discrepancy was present in his forearm bones, his sacro-iliac joints displayed ankylosis and he would have had genu valgum during life. Unlike Sk331, he was estimated to have been of roughly average height for the population (166.8 cm). Neither individual appears to have suffered from any tumors that progressed to malignancy^4^.

**Figure 1.**
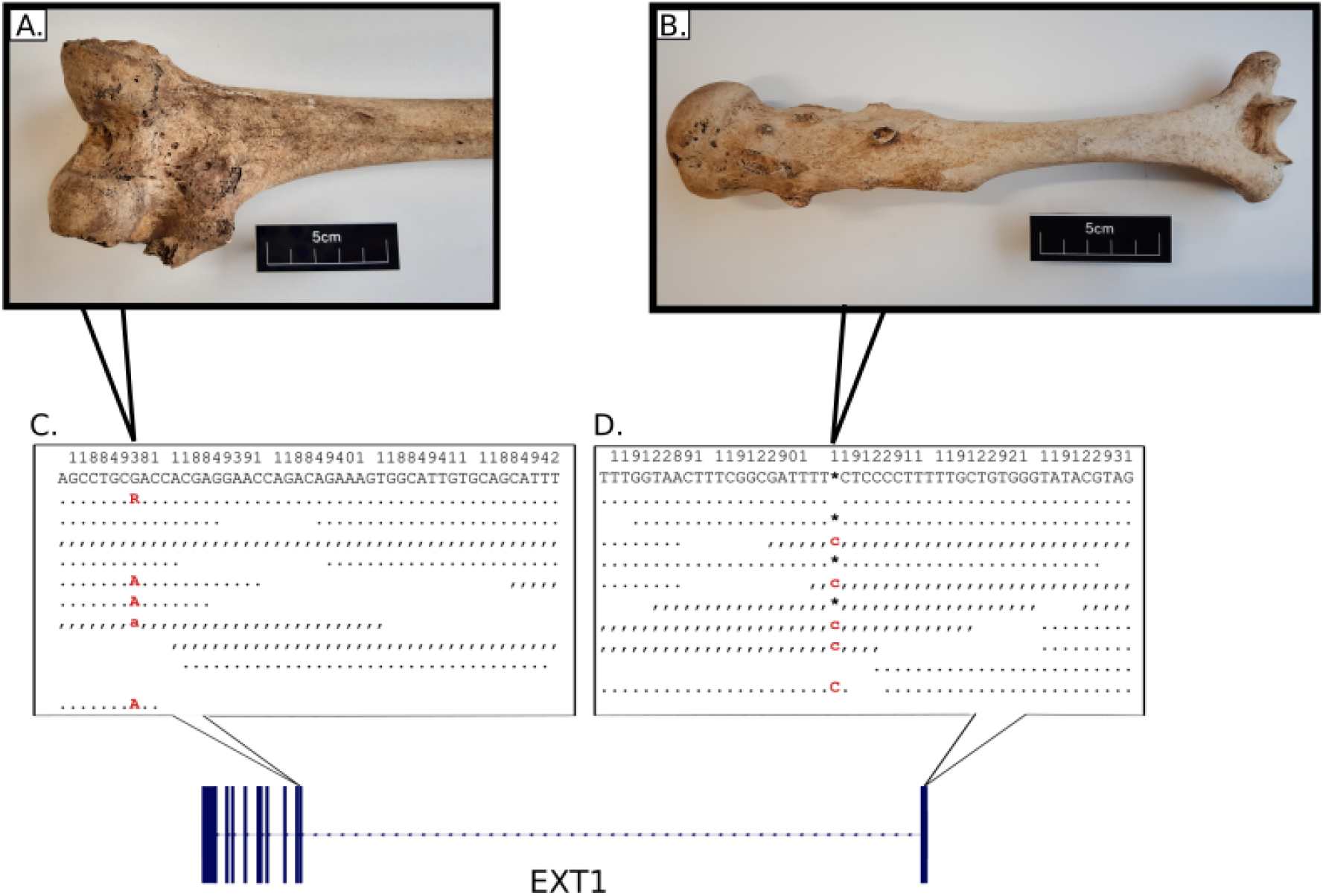
Examples of lesions and alignment of NGS data at putative disease variants. EXT1 gene structure with disease variants highlighted in red and bold typeface. First line: hs37d5 reference sequence. Second line: consensus for each individual. Dots: bases match reference on forward strand. Commas: bases match reference on reverse strand. R: consensus base is an A or G. A) Example of large sessile osteochondroma on the posterior surface of the distal end of Sk197’s left femur. B) Example of sessile and pedunculated osteochondromas on the posterior surface of Sk331’s left humerus. C) G>A transition in exon 2 of EXT1 in Sk197, resulting in Arginine to Cysteine mutation at the corresponding amino acid residue. D) C insertion in exon 1 of EXT1 in Sk331.

In order to identify likely causative mutations, we used an unbiased genome wide approach involving shotgun sequencing and ancient DNA protocols (see **Methods**). Petrous bone samples from each individual yielded intermediate levels of endogenous DNA preservation (Sk331: 12.2%; Sk197: 13.9%) and were sequenced to a mean depth of coverage of 4.2× and 5.1× respectively (see **Table 1** and **Table S2** for summary and sequencing statistics). Likely pathogenic mutations in *EXT1* were identified from an exome-wide scan in both individuals. All possible variants were initially filtered for quality (minimum allelic depth 3; maximum read depth twice the mean genomic coverage; minimum genotype quality 50) and predicted molecular impact using SnpEff and SnpSift (high or moderate impact)^7,8^. All qualifying variants were heterozygous calls. Therefore, only variants in genes with a high probability of loss of function intolerance (pLI > 90%) according to gnomADv2.1.1 were retained, in order to filter out variants that were unlikely to cause a pathological phenotype in a heterozygous state^9^. These variants were then assessed based on allele frequencies (<1% in gnomAD), *in silico* predicted pathogenicity (according to SIFT and Polyphen2) and publicly available data from ClinVar^9–11^. Only one mutation in each individual had the level of support required by the American College of Medical Genetics guidelines to classify them as pathogenic (**Figure 1; Table S2)**^12,13^.

**Table 1.**
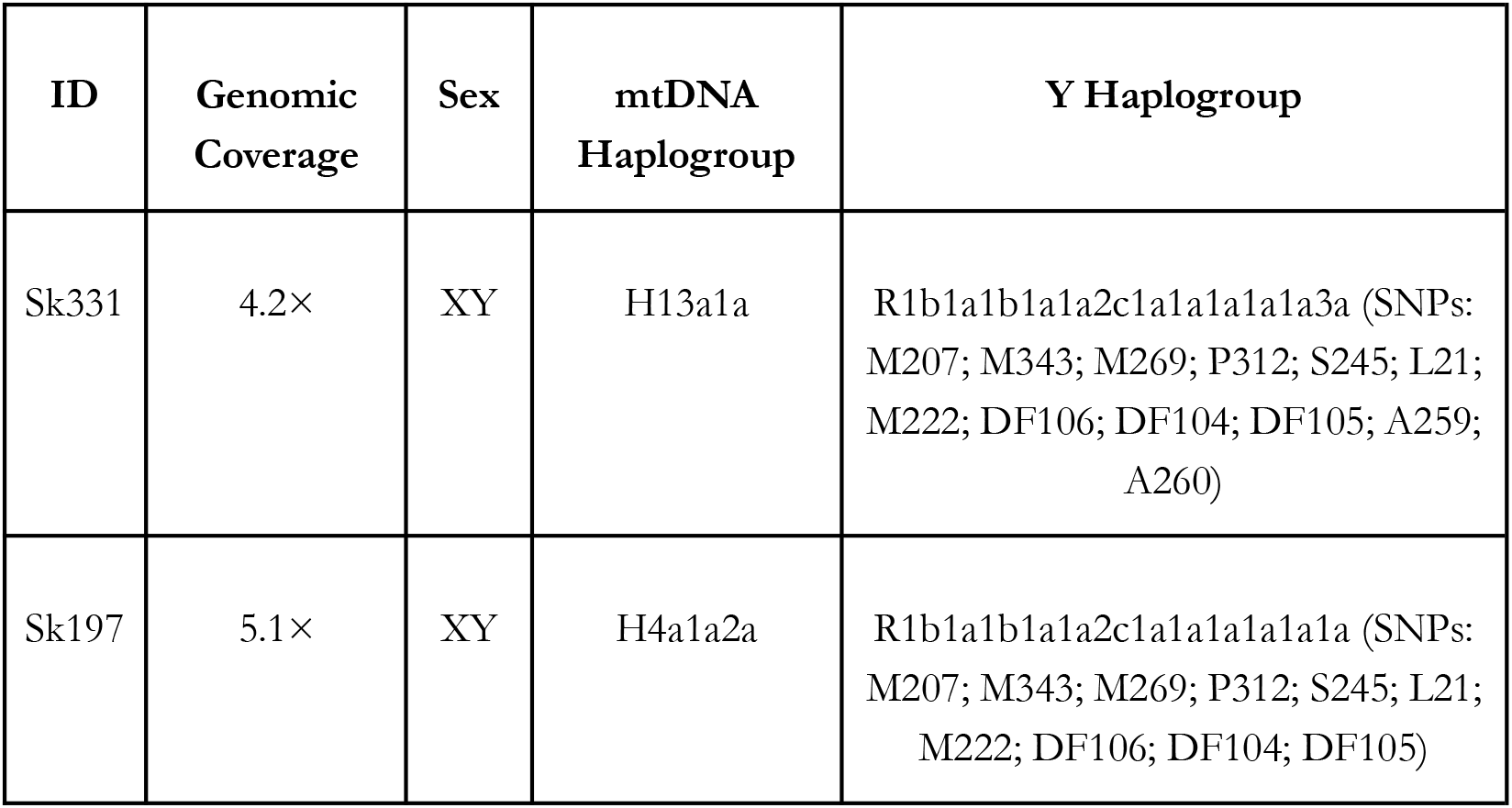
Sequencing Summary. Summary of coverage, sex and uniparental haplogroups assigned using MitoMaster and Phylotree 17 (mitochondrial) and ISOGG v15.58 (Y).

The predicted pathogenic mutation identified in Sk197 is a missense mutation in the second exon of *EXT1* (NC_000008.10:g.118849385G>A; NP000118:p.Arg340Cys) (**Figure 1c.**). This specific mutation has been identified in at least three patients^14–16^, and different missense mutations at the same amino acid residue have been identified as pathogenic (R340G,L,H: accessions VCV000988576, VCV000002495, VCV000265129), as have substitutions at nearby residues. Arginine to Cysteine is a non-conservative amino acid substitution, and at this position has been demonstrated to disrupt EXT1/EXT2 complex activity, consistent with what is known about disease mechanism^17,18^. Therefore, there are multiple lines of evidence supporting a pathogenic role for this mutation in Sk197: the same amino acid change as an established disease variant is observed; functional studies have shown a deleterious effect; this mutation is absent in population databases; computational evidence supports a damaging effect on the gene product; finally, the phenotype (MO) is highly specific for this gene.

Although the G>A mutation is not found in Sk331, a novel predicted pathogenic mutation is observed. This mutation is a C insertion within the first exon of *EXT1* (NC_000008.10:g.119122909_119122910insC; NP_000118:p.(Lys126Argfs*63)), resulting in a frameshift mutation and premature stop codon (**Figure 1d**.). This is a very severe mutation, resulting in a complete loss of the protein product of one copy of *EXT1* and is consistent with what is known about pathogenic mutations associated with this disease^19^. Although this mutation has not been observed in modern patients (as reported in ClinVar; date accessed 14-05-2021), there are 6 frameshift variants predicted to be pathogenic within 50bp of this site, and 3 nonsense SNVs predicted to be pathogenic in this region (**Table S4**). Therefore, this is a null mutation where loss of function is a known mechanism of disease, in a known disease gene, at a locus where there is modern clinical data supporting pathogenicity. Computational tools predict this variant to be deleterious. In sum, there is a high level of evidence to support the pathogenicity of this variant.

90% of cases of MO are caused by mutations in the exostosin genes *(EXT1* and *EXT2*), which are involved in heparan sulfate chain synthesis and assembly^20^. These chains interact with a wide range of signalling molecules, and deficiencies in these interactions lead to altered signalling pathways ^17^. Most of these mutations are classified as inactivating mutations; they result in a complete loss of the protein product from the copy of the gene carrying the mutation^19^. As *EXT1* and *EXT2* form a complex to carry out their molecular function, such mutations in either gene result in a reduction in the quantity of functional complex, and therefore a reduction in heparan sulfate chain production. Although this does have a physiological effect (for example, defects in lipid metabolism and clearance), it is insufficient for tumor formation^20^. MO is a dominant condition and it is thought that a complete somatic loss of EXT1/2 function is necessary for disease, which has been reported as being due to a loss of heterozygosity or aneuploidy^20,21^, although studies in mice have demonstrated that compound *EXT1/2* heterozygosity can mimic the human MO phenotype^22^.

To date only stature phenotypes in recent historical samples have led to successful identification of causal Mendelian lesions; none with a genome-wide approach. Two 18th and 19th century skeletons of extremely tall individuals had sufficiently intact DNA for targeted PCR of gigantism-associated loci^23,24,25^. A pathological achondroplasia mutation in the FGFR3 gene has also been identified in 180-year old remains^26^.

We also used projection principal components analysis with modern northwest European populations to test affinities of the two Ballyhanna genomes^27–29^. Both individuals fall at the overlap between Scottish and Irish samples, consistent with what we might expect for modern individuals from the northern part of Ireland^27^ (**Figure 2**).

**Figure 2.**
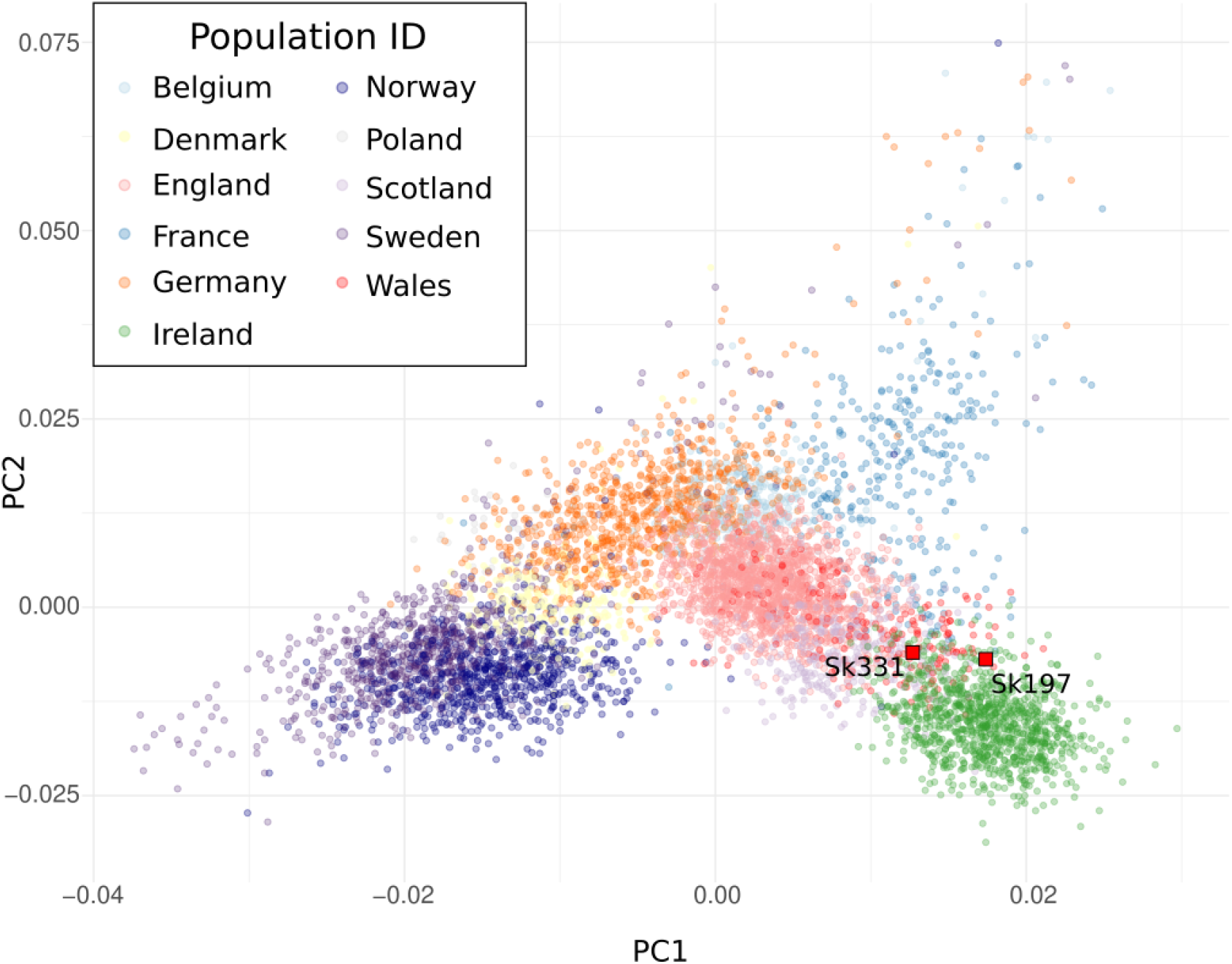
Principal Component Analysis of North-west European populations. Sk331 and Sk197 projected onto a plot of North-west European populations. Modern populations: Ireland (green); Scotland (lilac); Wales (red); England (pink); France (blue); Germany (orange); Belgium (pale blue); Norway (dark purple); Sweden (purple); Denmark (yellow); Poland (grey).

These men had different mitochondrial haplogroups but fell into the same clade of Y chromosome haplotypes, although Sk331 had a slightly more derived Y haplogroup (**Table 1**). Both grouped in the cluster R1b-M222, which is known to have its highest frequency in the same northwestern region in modern Ireland^30,31^. Interestingly, from modern examination of surnames, the Ó Domnaill Gaelic lords whose clan held power in this region would have been expected to display this haplotype^31^.

The haplotype background of each Ballyhanna pathogenic mutation was assessed using diploid data imputed using GLIMPSE and phased with a dataset of 72 imputed, published Iron Age and Medieval genomes using SHAPEITv2 (see **Table S5** for details)^32–36^. The region around *EXT1*, including 50kB up- and downstream were visualised using haplostrips^37^ (**Figure 3**). The Ballyhanna haplotypes do not cluster together, further supporting two independent MO-causing mutations in these individuals.

**Figure 3.**
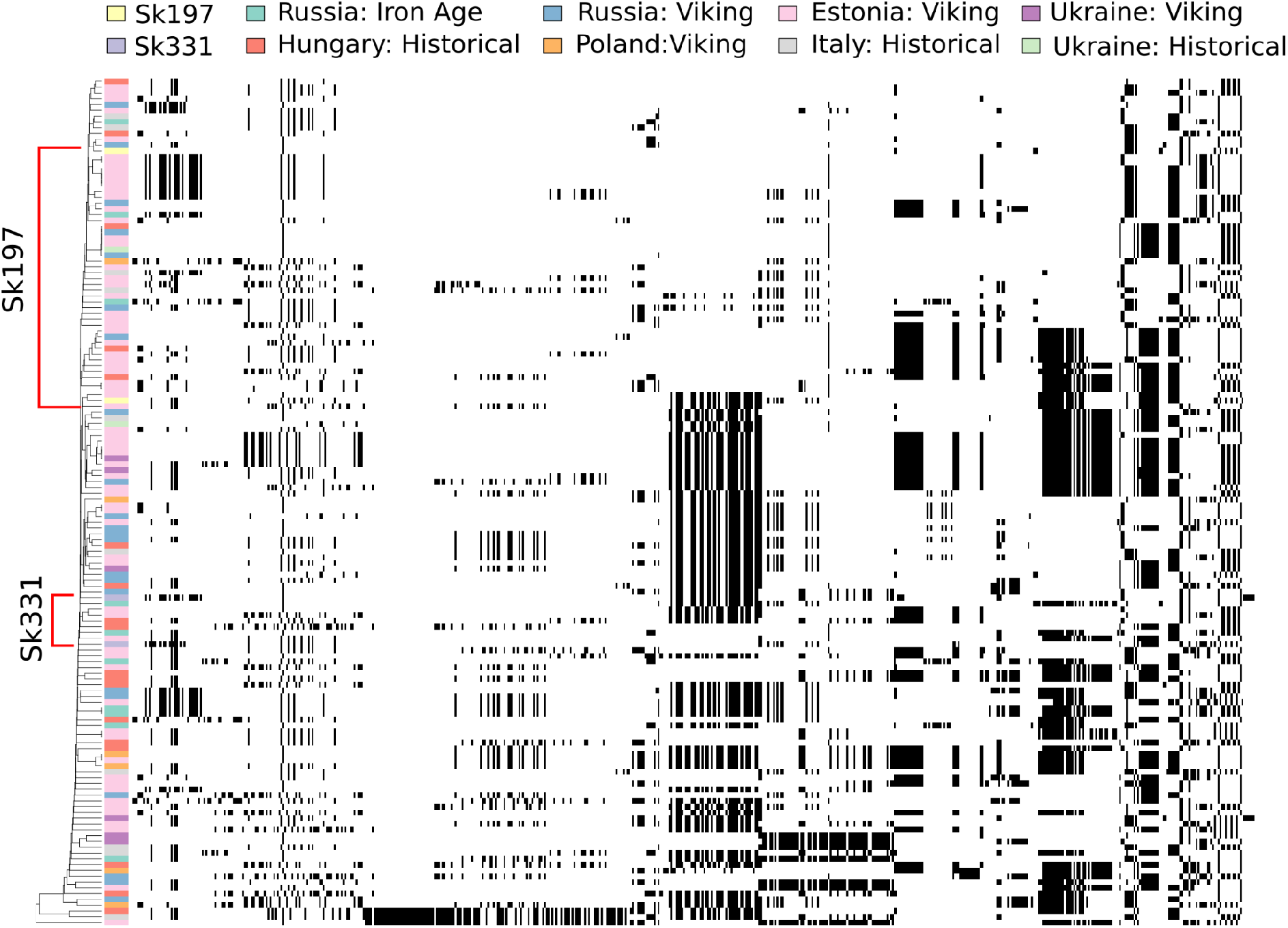
*EXT1* haplotype clustering in imputed ancient data. *EXT1* haplotypes were clustered using the tool haplostrips. Colours at the side of the plot indicate approximate geographic location and period of the individuals; Sk197 (pale yellow) and Sk331 (pale purple) are highlighted.

MO cases have been osteologically identified in the palaeopathological record from the Middle Bronze Age to the post-medieval period^6,38^. Ten of these cases are isolated examples, but four countries have multiple individuals with MO: Gotland (Sweden) (n=2: mother and unborn infant); Jordan (n=3, of which two are broadly contemporaneous), England (n=3) and Ireland (n=4, including the 2 probands here)^6^. The estimated incidence of this condition in modern individuals is approximately 1/50000, although higher incidences have been reported in the isolated populations of the Chamorros of Guam (1/1000) and the northern Ojibway community in Manitoba (13/1000)^39,40^. The *de novo* rate of mutation in this condition is low at 10%^41^, and this clustering in modern, restricted populations strongly suggests founder effects typical of severe dominant genetic diseases. Such would have been expected within the same graveyard in the north-western corner of Gaelic Medieval Ireland, and therefore it is striking that our analyses demonstrate these two rare cases arise from separate mutational events.

## Acknowledgements

We wish to thank Transport Infrastructure Ireland and Donegal County Council who funded the Ballyhanna Research Project and the National Museum of Ireland for granting the necessary licences for the study. This work was funded by the Science Foundation Ireland/Health Research Board/Wellcome Trust Biomedical Research Partnership Investigator Award No. 205072 to DGB, “Ancient Genomics and the Atlantic Burden” and IJ was supported by The Science Foundation Ireland Centre for Research Training in Genomics Data Science (18/CRT/6214). We acknowledge Trinseq for sequencing support and the DJEI/DES/SFI/HEA Irish Centre for High-End Computing (ICHEC) for the provision of computational facilities.

## Data and Code availability

Raw FASTQ and aligned BAM files are available from the European Nucleotide Archive (ENA) under accession number XXX.

## Supplemental information

Document S1

Tables S1-S5

## Methods

### Sampling and Sequencing Preparation

The petrous bones of two individuals with Multiple Osteochondromas from Ballyhanna, Co. Donegal (Excavation number 03E1384) were sampled for DNA extraction and shotgun sequencing (Licence to Alter Number 6861, National Museum of Ireland).

Bones were photographed before modification, and exposed to UV light for 30 minutes on either side to remove surface contaminants. The surface of each bone was cleaned using a drill bit, and a triangular section of the otic capsule was cut. Half was stored as a bone fragment, and the other half was pulverised in a Mixer Mill 400 (Retsch). Extracts were prepared from 0.1g of this powder. One extract of Sk331 was prepared as in Boessenkool et al (2017)^1^: bone powder was initially incubated in 0.5% bleach for 15 minutes to remove contaminants. After centrifuging for a few seconds at 13000 rpm, the bleach supernatant was discarded, and the remaining pellet was washed three times with UV water. The pellet was then incubated in UV-ed EDTA at 37°C for 30 minutes at 900rpm. This was then centrifuged at 13000 rpm for 10 minutes and the supernatant was removed and stored. The remaining pellet was incubated in UV-ed extraction buffer with proteinase K for 48 hours at 37°C and 900rpm. This was centrifuged at 13000 rpm for 10 minutes, and the supernatant was transferred to an Amicon Ultra-4 Centrifugal Filter unit 30kDa, diluted in 3mL 10 mM Tris-EDTA buffer and centrifuged at 5000 rpm until 100uL of solution remained. This volume was then added to a silica column (MinElute PCR purification kit, Qiagen, Hilden, Germany) and purified according to the manufacturer’s instructions. Sk197 was extracted following a similar protocol but with some modifications as in Dabney et al. (namely, shorter incubation with proteinase K (24hr), lower centrifugation speeds and the use of Zymo-Spin V columns (Zymo Research) rather than the Amicon columns)^2^. An additional extract from the initial EDTA wash was prepared for Sk197.

3 sequencing libraries were prepared from 16.25uL of each extract (total: 9 libraries). Each of these were USER-treated to mitigate the effects of post-mortem damage. Libraries were prepared as in Meyer and Kircher (2010), with modifications as in Gamba et al (2014)^3,4^. Libraries and controls were amplified using Accuprime Pfx (Life technology); indexing primers p5 and p7; 10x reaction buffer and 3uL of library/water with a total reaction volume of 25uL. The first libraries for each extract were amplified for 14 cycles for an initial screen. The remaining PCRs were amplified for 10-13 cycles for high coverage sequencing. PCRs were purified with the MinElute PCR purification kit, and the concentrations of these products were quantified using the Agilent Tapestation system with a D1000 screentape (Sk331) or a QuBit with the High Sensitivity dsDNA assay (Invitrogen) (Sk197). Samples were screened and sequenced to a high read depth on a Novaseq 6000 platform using paired-end 50bp reads.

### Alignment and Data Processing

The quality of demultiplexed fastq files was assessed using the FASTQC suite^5^. Adapter sequences were trimmed using AdapterRemoval v2.2.2 and paired end reads were collapsed ^6^. Trimmed reads were aligned to the hs37d5 reference genome using bwa aln with relaxed parameters and the seed disabled (-o 2 -n 0.01 -l 16500) ^7^. PCR and optical duplicates were removed using PicardTools v2.22.1 MarkDuplicates (http://broadinstitute.github.io/picard), and reads were filtered for mapping quality > 20 using samtools ^8^. Bam files were merged to sample level and indel realignment was performed using GATK ^9^. Reads were “softclipped” by reducing the quality score of the first and last two bases of each read to a PHRED score of two. Bam files were then fished for exact p7 index matches, as one base-pair mismatches had been allowed during demultiplexing.

Mitochondrial and X chromosome contamination estimates were used to assess authenticity of this data. Mitochondrial contamination was estimated using the frequency of heteroplasmic sites when aligned to the revised Cambridge Reference sequence, both including and excluding possible postmortem damage. X chromosome contamination levels were estimated using ANGSD as in Rasmussen et al. (2011) ^10^.

### Uniparental Haplogroups

Mitochondrial consensus sequences were estimated using samtools mpileup and vcfutils ^11^. Each site had a minimum read depth of 5 and base quality 30. Consensus fasta files were used as input to Mitomaster, which was used to estimate mitochondrial haplogroups ^12^.

Y haplogroups were investigated by calling representative SNPs of likely haplogroups from the International Society of Genetic Genealogy 2020 Y-DNA tree (v. 15.58) ^13^. Base calls across these loci were generated for both samples using GATK’s pileup tool, and loci with derived alleles were flagged. These were then manually assessed to call Y haplogroups.

### Pseudohaploid SNP calling

GATK was used to pileup SNPs of interest for pseudohaploid analyses from the final BAM files (see **Table S6** for details). These sites were filtered for minimum base quality 30, and one read was sampled at random to generate pseudohaploid calls.

### Modern Population Affinities

#### Outgroup f3 statistics

Shared drift with modern human populations was initially assessed using outgroup *f3* statistics using Admixtools ^14^. Pseudohaploid SNPs were called as above, samples were merged with the Human Origins dataset, and statistics of the form *f3*(Mbuti; Ballyhanna, Modern_Population) were calculated.

#### Principal Components Analysis

Principal components were calculated on a dataset of modern Northwestern Europeans (including Irish individuals) ^15–17^, and Sk331 and 197 were projected onto this using smartpca v8000 with shrinkmode:YES ^18,19^. No outlier iterations were performed, and SNPs in LD were eliminated with an r^2^ threshold > 0.2.

### Exome Scan

A scan to search for possible dominant acting variants was developed. Bcftools v1.10.2 was used to call possible variant sites in autosomal exons from bam files aligned to hs37d5 ^11^, and calls were filtered for quality using the command

> “bcftools mpileup -R ${bed} --count-orphans --annotate FORMAT/AD,INFO/AD --ignore-RG --fasta-ref ${ref} ${bamfile} -Ou | bcftools call -mv --format-fields GQ,GP-Ou | bcftools filter -s LowQual -Oz -o ${id}.exome.unsorted.vcf.gz”.

Co-ordinates of autosomal exons were downloaded from the UCSC table browser tool on 15-09-2020 in BED format using the NCBI RefSeq database, including 2bp up- and downstream of the coding sequences ^20^.

SnpEff and SnpSift were used to annotate these possible variants in terms of molecular impact on canonical transcripts ^21,22^. Possible variants were filtered for minimum allelic depth 3 and maximum read depth less than twice the mean genomic coverage. A minimum genotyping quality of 50 was used. Sites were then filtered for high or moderate molecular impact, and variants in genes with a high probability of loss of function intolerance (pLI > 90%) according to gnomADv2.1.1 were retained ^23^. These variants were filtered for low (<1%) allele frequency in gnomad and predicted pathogenic effect by both SIFT and Polyphen2 (SNPs) or just SIFT (indels) ^24,25^. The ACMG classification guidelines were used to assess support for the candidate pathogenic variants ^26^.

### Imputation

Autosomal imputation was performed on both individuals using GLIMPSE ^27^. Genotype likelihoods for autosomal biallelic SNPs in the 1000 genomes project phase 3 panel were called for each individual using bcftools v1.10.2 as in the GLIMPSE documentation ^11,27^. Individuals were merged on a per-chromosome basis and imputed in chunks of 2Mb with a 200kB buffer region, using the Phase 3 panel of the 1000 genomes project as a reference dataset^28^. A genotype probability filter of 99% was used for all downstream analyses of this data.

### EXT1 Haplotypes

Relationships between the haplotypes at the disease gene *(EXT1)* in affected individuals was investigated in the context of a larger imputed dataset of published Iron Age and Historical samples^29–32^. This data was phased using SHAPEITv2 with the 1000 Genomes Project Phase 3 reference panel^28,33^. The region spanned by *EXT1* exons and 50kB up- and down-stream were visualised using the tool haplostrips ^34^.

### Radiocarbon Dating

Radiocarbon dating was undertaken on two rib samples. The dates were repeated to ensure accuracy.

| Lab Code | Sample/Context | Years BP | Calibrated Date Ranges |
| --- | --- | --- | --- |
| UBA-11442 | Rib from Sk197 | 1223 +/- 27 | AD 708-877 (1 $\sigma$);<br>AD 689-885 (2 $\sigma$) |
| UBA-11442 Repeat | | 1162 +/- 24 | AD 775-950 (1 $\sigma$);<br>AD 774-975 (2 $\sigma$) |
| UBA-11443 | Rib from Sk331 | 856 +/- 24 | AD 1175-1221 (1 $\sigma$);<br>AD 1158-1260 (2 $\sigma$) |
| UBA-11443 Repeat | | 928 +/- 40 | AD 1042-1162 (1 $\sigma$);<br>AD 1031-1211(2 $\sigma$) |

Calibration: Reimer et al, 2020 <sup>35</sup>

## Supplementary figures

**Figure S1.**
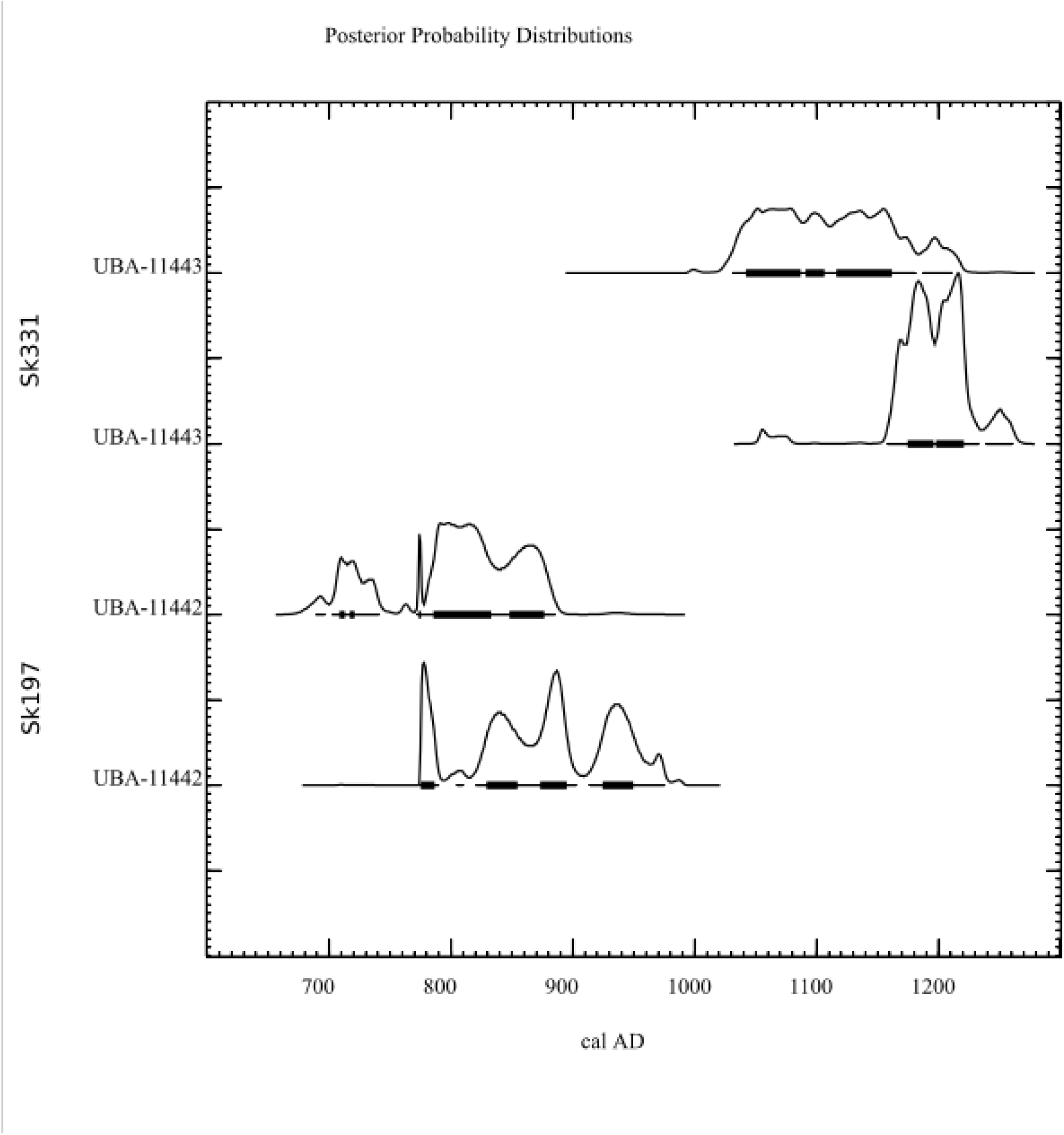
Radiocarbon dates for Sk197 and Sk331. Posterior distribution of estimated age is displayed, where the x-axis is in calendar years AD. UBA-11442 is Sk197 and UBA-11443 is Sk331.

**Figure S2.**
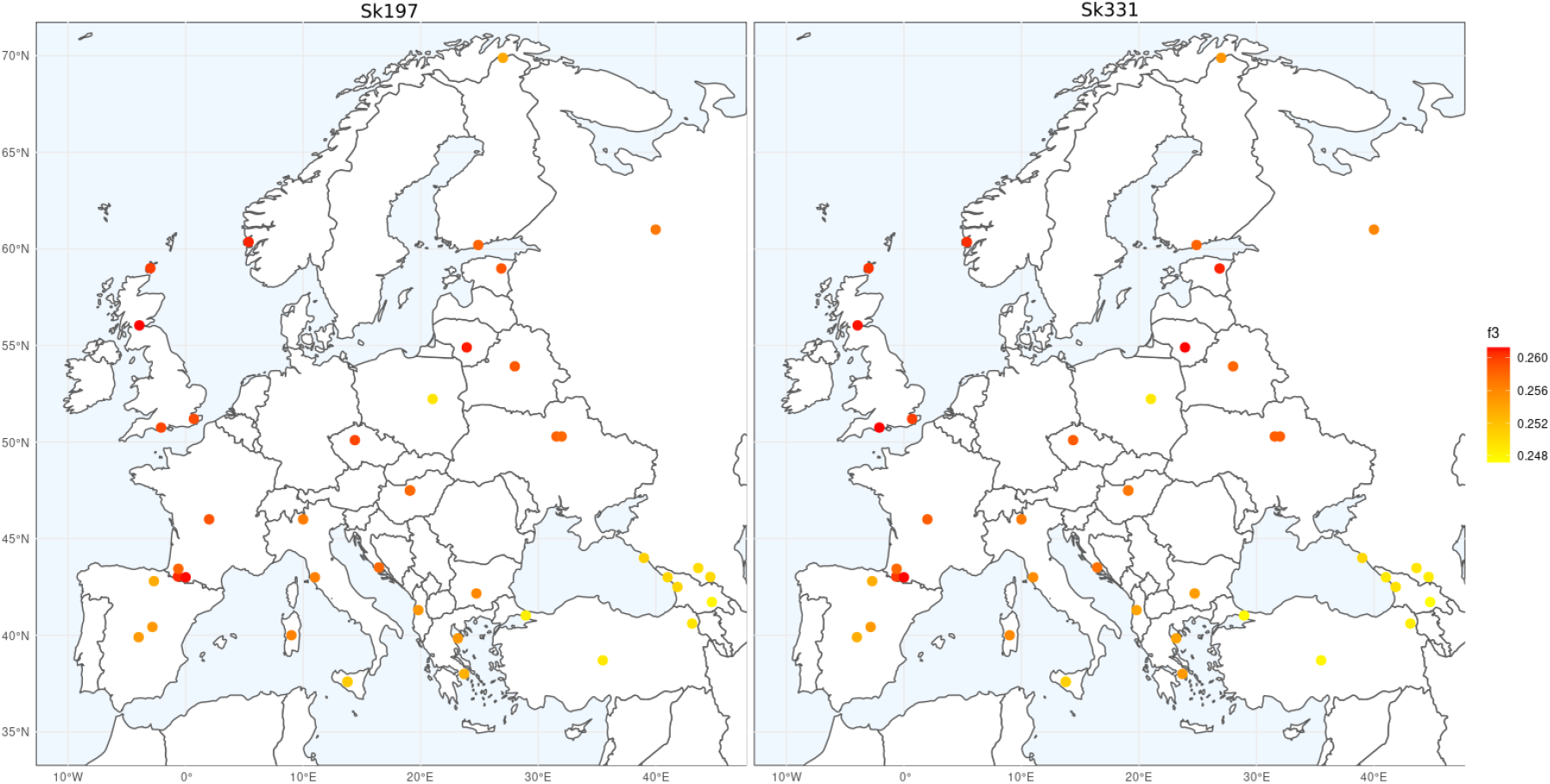
Outgroup *f3* statistics with European Populations. Shared drift with modern European populations assessed using the statistic *f3*(Mbuti; Ballyhanna, Modern_Population). Higher values are closer to red and indicate more shared drift.

